## Supplementary Materials for "Dopamine challenge reduces mental state attribution accuracy"

Appendix 1. Model parameters

**Supplementary Table S1.1** *Model parameters for model 1.1*

| **Population-level effects** | **Estimate** | **Error** | **95% CrI (lower)** | **95% CrI (upper)** |
|---|---|---|---|---|
| *Intercept* | 4.27 | 0.27 | 3.73 | 4.81 |
| *HAL vs PLA* | -0.56 | 0.19 | -0.94 | -0.19 |
| **Group-level effects** | **Estimate (SD)** | **Error** | **95% CrI (lower)** | **95% CrI (upper)** |
| *Subject ID (Intercept)* | 1.13 | 0.16 | 0.86 | 1.47 |
| *Subject ID (drug)* | 0.68 | 0.22 | 0.20 | 1.12 |
| *Animation ID (Intercept)* | 2.20 | 0.15 | 1.92 | 2.51 |

*Note.* Model formula: accuracy ~ drug + (1 + drug || subject ID) + (1 | animation ID).

**Supplementary Table S1.2**. *Model parameters for model 1.2*

| **Population-level effects** | **Estimate** | **Error** | **95% CrI (lower)** | **95% CrI (upper)** |
|---|---|---|---|---|
| *Intercept* | 5.36 | 0.31 | 4.75 | 5.95 |
| *HAL vs PLA* | -0.66 | 0.23 | -1.12 | -0.20 |
| *Mental vs non-mental* | -2.50 | 0.37 | -3.22 | -1.78 |
| *HAL vs PLA, mental vs non-mental* | 0.20 | 0.28 | -0.34 | 0.74 |
| **Group-level effects** | **Estimate (SD)** | **Error** | **95% CrI (lower)** | **95% CrI (upper)** |
| *Subject ID (Intercept)* | 1.12 | 0.16 | 0.85 | 1.47 |
| *Subject ID (drug)* | 0.69 | 0.22 | 0.23 | 1.11 |
| *Animation ID (Intercept)* | 1.84 | 0.13 | 1.60 | 2.13 |

*Note.* Model formula: accuracy ~ drug * mental state + (1 + drug || subject ID) + (1 | animation ID).

**Supplementary Table S1.3** *Model parameters for model 1.3*

| **Population-level effects** | **Estimate** | **Error** | **95% CrI (lower)** | **95% CrI (upper)** |
|---|---|---|---|---|
| *Intercept* | 4.21 | 0.36 | 3.50 | 4.92 |
| *HAL vs PLA* | -0.52 | 0.28 | -1.08 | 0.03 |
| *Drug day 1 vs 2* | 0.12 | 0.42 | -0.71 | 0.93 |
| *HAL vs PLA, Drug day 1 vs 2* | -0.06 | 0.38 | -0.81 | 0.68 |

| **Group-level effects** | **Estimate (SD)** | **Error** | **95% CrI (lower)** | **95% CrI (upper)** |
|---|---|---|---|---|
| *Subject ID (Intercept)* | 1.14 | 0.16 | 0.87 | 1.49 |
| *Subject ID (drug)* | 0.72 | 0.22 | 0.27 | 1.15 |
| *Animation ID (Intercept)* | 2.20 | 0.15 | 1.92 | 2.52 |

*Note.* Model formula: accuracy ~ drug * drug day + (1 + drug || subject ID) + (1 | animation ID).

**Supplementary Table S2.1**. *Model parameters for model 2.1*

| **Population-level effects** | **Estimate** | **Error** | **95% CrI (lower)** | **95% CrI (upper)** |
|---|---|---|---|---|
| *Intercept* | 5.39 | 0.30 | 4.81 | 5.99 |
| *HAL vs PLA* | -0.69 | 0.23 | -1.14 | -0.22 |
| *Mental vs non-mental* | -2.71 | 0.37 | -3.47 | -2.00 |
| *Jerk difference* | -0.11 | 0.13 | -0.36 | 0.14 |
| *HAL vs PLA, mental vs non-mental* | 0.44 | 0.31 | -0.17 | 1.06 |
| *HAL vs PLA, jerk difference* | 0.06 | 0.17 | -0.26 | 0.39 |
| *Mental vs non-mental, jerk difference* | -0.54 | 0.28 | -1.09 | -0.00 |
| *HAL vs PLA, mental vs non-mental, jerk difference* | 0.68 | 0.41 | -0.11 | 1.48 |

| **Group-level effects** | **Estimate (SD)** | **Error** | **95% CrI (lower)** | **95% CrI (upper)** |
|---|---|---|---|---|
| *Subject ID (Intercept)* | 1.09 | 0.15 | 0.82 | 1.43 |
| *Subject ID (drug)* | 0.66 | 0.22 | 0.20 | 1.08 |
| *Animation ID (Intercept)* | 1.81 | 0.14 | 1.56 | 2.09 |

*Note.* Model formula: accuracy ~ drug * mental state * jerk difference + (1 + drug || subject ID) + (1 | animation ID).

**Supplementary Table S2.2**. *Model parameters for model 2.2 (PLA only)*

| **Population-level effects** | **Estimate** | **Error** | **95% CrI (lower)** | **95% CrI (upper)** |
|---|---|---|---|---|
| *Intercept* | 5.39 | 0.30 | 4.80 | 5.98 |
| *Jerk difference* | -0.13 | 0.14 | -0.41 | 0.14 |
| *Mental vs non-mental* | -2.76 | 0.37 | -3.47 | -2.03 |

| | | | | |
|---|---|---|---|---|
| *Jerk difference, mental vs non-mental* | -0.70 | 0.31 | -1.32 | -0.09 |

| **Group-level effects** | **Estimate (SD)** | **Error** | **95% CrI (lower)** | **95% CrI (upper)** |
|---|---|---|---|---|
| *Subject ID (Intercept)* | 1.07 | 0.18 | 0.76 | 1.48 |
| *Animation ID (Intercept)* | 1.72 | 0.16 | 1.42 | 2.05 |

*Note.* Model formula: accuracy ~ jerk difference * mental state + (1 | subject ID) + (1 | animation ID).

**Supplementary Table S2.3**. *Model parameters for model 2.3 (HAL only)*

| **Population-level effects** | **Estimate** | **Error** | **95% CrI (lower)** | **95% CrI (upper)** |
|---|---|---|---|---|
| *Intercept* | 4.74 | 0.32 | 4.10 | 5.36 |
| *Jerk difference* | -0.10 | 0.14 | -0.37 | 0.18 |
| *Mental vs non-mental* | -2.26 | 0.38 | -2.99 | -1.52 |
| *Jerk difference, mental vs non-mental* | 0.02 | 0.36 | -0.69 | 0.73 |

| **Group-level effects** | **Estimate (SD)** | **Error** | **95% CrI (lower)** | **95% CrI (upper)** |
|---|---|---|---|---|
| *Subject ID (Intercept)* | 1.36 | 0.19 | 1.02 | 1.78 |
| *Animation ID (Intercept)* | 1.74 | 0.16 | 1.46 | 2.07 |

*Note.* Model formula: accuracy ~ jerk difference * mental state + (1 | subject ID) + (1 | animation ID).

**Supplementary Table 3**. *Model parameters for model 3*

| **Population-level effects** | **Estimate** | **Error** | **95% CrI (lower)** | **95% CrI (upper)** |
|---|---|---|---|---|
| *Intercept* | 5.39 | 0.30 | 4.79 | 5.98 |
| *HAL vs PLA* | -0.74 | 0.25 | -1.23 | -0.26 |
| *Mental vs non-mental* | -2.75 | 0.37 | -3.47 | -2.04 |
| *PLA jerk difference* | -0.16 | 0.13 | -0.42 | 0.10 |
| *HAL vs PLA, mental vs non-mental* | 0.35 | 0.31 | -0.26 | 0.96 |
| *HAL vs PLA, PLA jerk difference* | 0.16 | 0.17 | -0.18 | 0.50 |
| *Mental vs non-mental, PLA jerk difference* | -0.55 | 0.28 | -1.10 | 0.00 |
| *HAL vs PLA, mental vs non-mental, PLA jerk difference* | 0.19 | 0.38 | -0.56 | 0.94 |

| **Group-level effects** | **Estimate (SD)** | **Error** | **95% CrI (lower)** | **95% CrI (upper)** |
|---|---|---|---|---|
| *Subject ID (Intercept)* | 1.15 | 0.18 | 0.85 | 1.53 |
| *Subject ID (drug)* | 0.74 | 0.23 | 0.25 | 1.21 |
| *Animation ID (Intercept)* | 1.78 | 0.14 | 1.53 | 2.06 |

*Note.* Model formula: accuracy ~ drug * mental state * PLA jerk difference + (1 + drug || subject ID) + (1 | animation ID).

**Supplementary Table 4.1**. *Model parameters for model 4.1*

| **Population-level effects** | **Estimate** | **Error** | **95% CrI (lower)** | **95% CrI (upper)** |
|---|---|---|---|---|
| *Intercept* | -0.05 | 0.02 | -0.10 | -0.00 |
| *ER change* | -0.02 | 0.02 | -0.07 | 0.03 |
| *Mental vs non-mental* | -0.02 | 0.03 | -0.08 | 0.05 |
| *WM change* | -0.00 | 0.00 | -0.01 | 0.01 |
| *ER change, mental vs non-mental* | 0.06 | 0.03 | -0.00 | 0.13 |
| *WM change, mental vs non-mental* | -0.01 | 0.00 | -0.02 | 0.00 |

*Note.* Model formula: accuracy change | trunc(lb = -1, ub = 1) ~ emotion change * mental state + WM change * mental state. ER change = emotion recognition change; WM change = working memory change.

Model 4.1 was modelled as a simple linear model as model comparison (using the function loo_compare of the LOO package[1]) with the same model including a random intercept for subject ID (model 4.1.rand) did not favour the random effects model:

**Supplementary Table 4.2**. *Leave-one-out (loo) cross-comparison of models 4.1 and 4.1.rand*

| | elpd_diff | se_diff |
|---|---|---|
| **Model 4.1** | 0.0 | 0.0 |
| **Model 4.1.rand** | -0.7 | 0.8 |

*Note*. Elpd_diff = Bayesian LOO estimate of the expected log pointwise predictive density (see[1]); se_diff = standard error of elpd_diff.

Furthermore, because the response variable 'accuracy change' is not continuous, we re-ran model 4.1 specifying a cumulative instead of the student's t distribution (model 4.1.cum) and compared this to a discretised (using piecewise constant approximation of the continuous function) continuous model. Model comparison clearly favoured the student's t over the cumulative distribution family:

**Supplementary Table 4.3**. *Leave-one-out (loo) cross-comparison of models 4.1 and 4.1.cum*

| | elpd_diff | se_diff |
|---|---|---|
| **Model 4.1** | 0.0 | 0.0 |
| **Model 4.1.cum** | -33.9 | 6.1 |

*Note*. Elpd_diff = Bayesian LOO estimate of the expected log pointwise predictive density (see[1]); se_diff = standard error of elpd_diff.

**Supplementary Table 4.3**. *Model parameters for model 4.2*

| Population-level effects | Estimate | Error | 95% CrI (lower) | 95% CrI (upper) |
|---|---|---|---|---|
| *Intercept* | -0.05 | 0.02 | -0.09 | -0.00 |
| *ER change* | -0.02 | 0.02 | -0.07 | 0.03 |
| *Mental vs non-mental* | -0.01 | 0.03 | -0.08 | 0.06 |
| *ER change, mental vs non-mental* | 0.07 | 0.03 | -0.00 | 0.13 |

*Note.* Model formula: accuracy change | trunc(lb = -1, ub = 1) ~ ER change * mental state. ER change = emotion recognition change.

Appendix 2. Supplementary Results

### 2.1. Modelling the bimodality of the response

Due to bimodality of the response variable (see Fig. S1A), model 1.2 was re-run (Model 5, without fitting a random slope for animation ID due to convergence issues), this time modelling the response variable as a mixture of two gaussian distributions. Figure S1B shows that this captured the response structure reasonably well (note that a mixture of two skew-normal distributions would have provided an even better posterior fit but was not chosen due to convergence problems and to avoid overfitting). This approach resulted in a set of parameters for each of the two modelled distribution components. The model revealed a main effect of drug for the second distribution component ($E\mu_{2,HALvsPLA,non-mental}$ = -0.50, CrI = [-0.76, -0.25]; $E\mu_{2,HALvsPLA,mentalVSnon-mental}$ = 0.00, CrI = [-0.38, 0.39]), comprising all accuracy values above the distribution intersection point of 3.01 (determined using the

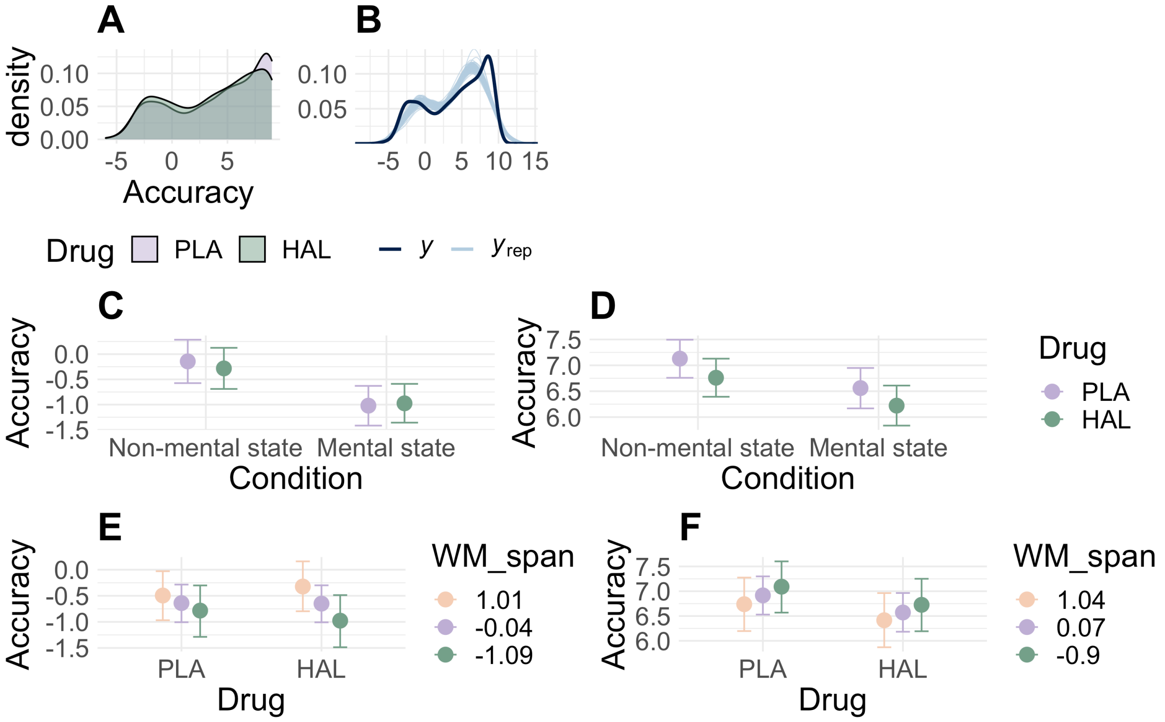


**Figure S1.** PLA = placebo trials, HAL = haloperidol trials. A) Probability density plot of the response variable accuracy. B) Posterior probability distribution of model 1.2. Y = response distribution, $y_{rep}$ = 100 draws from posterior samples. C-F) conditional effects plots[1] for simple gaussian models fit to each distribution component individually, C-D) depicting the interaction term of drug and mental state; E-F) depicting the interaction term of drug and baseline WM. C = distribution component 1 comprising accuracy values < 3.01, D = distribution component 2 with accuracy values > 3.01. E = distribution component 1 comprising accuracy values < 3.01, E = distribution component 2 with accuracy values > 3.01.

---

[1] Plots made using the 'conditional_effects()' function of the brms package (2 Bürkner, P.-C. brms: An R Package for Bayesian Multilevel Models Using Stan. *2017* **80**, 28, doi:10.18637/jss.v080.i01 (2017). Conditional effects plots display main effects and interactions of predictors on the response conditional on all predictors in the model.

uniroot.all function of the rootSolve package[3]). As in model 1.2, there was no interaction of drug and mental state for either of the distribution components (see Figure S1C-D; Supplementary Table 5.1). Thus, after taking into account the bimodality of the response variable, the model shows the same negative effect of drug on accuracy for mental- and non-mental state animations, whereby this model highlights that haloperidol affected accuracy particularly for animations that were initially decoded with higher accuracy. Importantly, we show below that dopamine challenge also affected performance for the more difficult animations (first distribution component), but here likely via different mechanisms. Finally, for both distribution components, the model revealed the same main effect of mental state as model 1.2 ($E\mu_{1,mentalVSnon-mental}$ = -1.17, CrI = [-1.63, -0.70]; $E\mu_{2,mentalVSnon-mental}$ = -0.81, CrI = [-1.09, -0.53]).

**Supplementary Table 5**. *Model parameters for model 5*

| **Population-level effects** | **Estimate** | **Error** | **95% CrI (lower)** | **95% CrI (upper)** |
|---|---|---|---|---|
| *Intercept$_1$* | -0.42 | 0.22 | -0.86 | 0.01 |
| *Intercept$_2$* | 6.87 | 0.17 | 6.54 | 7.19 |
| *HAL vs PLA$_1$* | -0.25 | 0.27 | -0.76 | 0.28 |
| *Mental vs non-mental$_1$* | -1.17 | 0.24 | -1.63 | -0.70 |
| *HAL vs PLA, mental vs non-mental$_1$* | 0.29 | 0.32 | -0.35 | 0.92 |
| *HAL vs PLA$_2$* | -0.50 | 0.13 | -0.76 | -0.25 |
| *Mental vs non-mental$_2$* | -0.81 | 0.14 | -1.09 | -0.53 |
| *HAL vs PLA, mental vs non-mental$_2$* | 0.00 | 0.20 | -0.38 | 0.39 |
| **Group-level effects** | **Estimate (SD)** | **Error** | **95% CrI (lower)** | **95% CrI (upper)** |
| *Subject ID (Intercept$_1$)* | 0.82 | 0.15 | 0.56 | 1.15 |
| *Subject ID (Intercept$_2$)* | 1.35 | 0.17 | 1.07 | 1.72 |

*Note.* Model formula: accuracy ~ drug * mental state + (1 | subject ID). Response modelled as a mixture of two gaussian distributions.

Based on previous evidence indicating that working memory span reliably predicts individual dopamine synthesis capacity[4,5] and drug effects of working memory span on performance in other socio-cognitive[6] and cognitive[7-11] domains, we used individual baseline working memory span as a proxy for baseline dopamine function, to account for interindividual differences in drug responsivity. For this, working memory task accuracy was calculated as the percentage of correct responses of all placebo trials. Participants were then divided into groups of low and high baseline working memory span by performing a median split on working memory span scores from the placebo day (indexing low and high dopamine synthesis capacity, respectively; 18 subjects in the low, 20 in the high working memory group).

A second mixture model (Model 6.1, see Supplementary Table 5.2) was fit to drug and *baseline working memory* (low, high; effects coded), as well as their interaction, predicting accuracy. This second model revealed an interaction between baseline working memory and drug for both distribution components, indicating a stronger decrease in the low, relative to the high WM group ($E\mu_{1,HALvsPLA,lowWM}$ = -0.34, CrI = [-0.68, 0.01], $E\mu_{1,HALvsPLA,highWM}$ = 0.34, CrI = [-0.01, 0.68]; $E\mu_{2,HALvsPLA,lowWM}$ = -0.30, CrI = [-0.51, -0.10], $E\mu_{2,HALvsPLA,highWM}$ = 0.30, CrI = [0.10, 0.51]). Two separate post-hoc models for low and high WM groups confirm a negative negative effect of drug on accuracy for both distribution components in low WM individuals (Model 6.2: $E\mu_{1,HALvsPLA}$ = -0.37, CrI = [-0.81, 0.06], $E\mu_{2,HALvsPLA}$ = -0.83, CrI = [-1.16, -0.50]), whereas the high WM group exhibited a negative drug effect for the second, but not the first distribution component (Model 6.3: $E\mu_{1,HALvsPLA}$ = 0.21, CrI = [-0.36, 0.77], $E\mu_{2,HALvsPLA}$ = -0.21, CrI = [-0.46, 0.04]; see Fig S1E-F). Thus, for those animations classified with accuracy values below 3.01, the effect of haloperidol on accuracy depended on individuals' baseline working memory capacity, whereby only individuals with low WM, and thus estimated low striatal dopamine synthesis capacity, showed decreased accuracy as a response to the drug. In contrast, both WM groups exhibited a decrease in accuracy after haloperidol for the

**Supplementary Table 6.1**. *Model parameters for model 6.1*

| Population-level effects | Estimate | Error | 95% CrI (lower) | 95% CrI (upper) |
|---|---|---|---|---|
| *Intercept₁* | -1.08 | 0.16 | -1.39 | -0.78 |
| *Intercept₂* | 6.49 | 0.17 | 6.16 | 6.82 |
| *HAL vs PLA₁* | -0.02 | 0.17 | -0.36 | 0.32 |
| *Low_WM₁* | -0.04 | 0.18 | -0.39 | 0.30 |
| *High_WM₁* | 0.04 | 0.18 | -0.30 | 0.39 |
| *HAL vs PLA, low_WM₁* | -0.34 | 0.18 | -0.68 | 0.01 |
| *HAL vs PLA, high_WM₁* | 0.34 | 0.18 | 0.01 | -0.68 |
| *HAL vs PLA₂* | -0.52 | 0.11 | -0.73 | -0.31 |
| *Low_WM₂* | 0.30 | 0.26 | -0.21 | 0.80 |
| *High_WM₂* | -0.30 | 0.26 | -0.21 | 0.80 |
| *HAL vs PLA, low_WM₂* | -0.30 | 0.11 | -0.51 | -0.10 |
| *HAL vs PLA, high_WM₂* | 0.30 | 0.11 | 0.10 | 0.51 |

| Group-level effects | Estimate (SD) | Error | 95% CrI (lower) | 95% CrI (upper) |
|---|---|---|---|---|
| *Subject ID (Intercept₁)* | 0.76 | 0.15 | 0.50 | 1.07 |
| *Subject ID (Intercept₂)* | 1.51 | 0.20 | 1.18 | 1.95 |

*Note.* Model formula: accuracy ~ drug * WM + (1 | subject ID). WM = working memory; response modelled as a mixture of two gaussian distributions.

animations with accuracy values above 3.01, albeit with a somewhat stronger negative effect in the low WM group.

**Supplementary Table 6.2**. *Model parameters for model 6.2 (post-hoc model - low WM)*

| **Population-level effects** | **Estimate** | **Error** | **95% CrI (lower)** | **95% CrI (upper)** |
|---|---|---|---|---|
| *Intercept$_1$* | -1.24 | 0.18 | -1.60 | -0.88 |
| *Intercept$_2$* | 6.53 | 0.20 | 6.14 | 6.91 |
| *HAL vs PLA$_1$* | -0.37 | 0.22 | -0.81 | 0.06 |
| *HAL vs PLA$_2$* | -0.83 | 0.17 | -1.16 | -0.50 |
| **Group-level effects** | **Estimate (SD)** | **Error** | **95% CrI (lower)** | **95% CrI (upper)** |
| *Subject ID (Intercept$_1$)* | 0.80 | 0.20 | 0.48 | 1.24 |
| *Subject ID (Intercept$_2$)* | 1.69 | 0.31 | 1.19 | 2.41 |

*Note.* Model formula: accuracy ~ drug + (1 | subject ID). Response modelled as a mixture of two gaussian distributions.

**Supplementary Table 6.3**. *Model parameters for model 6.3 (post-hoc model - high WM)*

| **Population-level effects** | **Estimate** | **Error** | **95% CrI (lower)** | **95% CrI (upper)** |
|---|---|---|---|---|
| *Intercept$_1$* | -1.20 | 0.23 | -1.66 | -0.76 |
| *Intercept$_2$* | 6.24 | 0.19 | 5.87 | 6.60 |
| *HAL vs PLA$_1$* | 0.21 | 0.29 | -0.36 | 0.77 |
| *HAL vs PLA$_2$* | -0.21 | 0.13 | -0.46 | 0.04 |
| **Group-level effects** | **Estimate (SD)** | **Error** | **95% CrI (lower)** | **95% CrI (upper)** |
| *Subject ID (Intercept$_1$)* | 1.04 | 0.33 | 0.48 | 1.79 |
| *Subject ID (Intercept$_2$)* | 1.54 | 0.30 | 1.09 | 2.24 |

*Note.* Model formula: accuracy ~ drug + (1 | subject ID). Response modelled as a mixture of two gaussian distributions.

### 2.2. Dopamine challenge reduced walking speed in individuals with low estimated dopamine synthesis capacity[2]

A Bayesian mixed effects model (Model 7) of drug (PLA, HAL; dummy coded) and WM group (low, high; effects coded) predicting walking speed revealed a negative main effect of drug ($E\mu_{PLAvsHAL}$ = -0.04, CrI = [-0.08, 0.01], P($E\mu_{PLAvsHAL}$) < 0 = 0.95), indicating that, overall,

[2] Results published in our previous paper: Dopaminergic Modulation of Dynamic Emotion Perception (Schuster et al., 2022, *JNeurosci)*.

haloperidol tended to reduce walking speed. In addition, there was a main effect of WM group ($E\mu_{lowWM}$ = -0.04, CrI = [-0.09, 0.01], $E\mu_{highWM}$ = 0.04, CrI = [-0.01, 0.09]), demonstrating that under placebo, the low WM group exhibited a slower walking pace relative to high WM individuals (low WM: mean [M] = 1.05 m/s, high WM: M = 1.13 m/s). There further was an interaction between drug and WM group ($E\mu_{PLAvsHAL,lowWM}$ = -0.04, CrI = [-0.09, 0.00], $E\mu_{PLAvsHAL,highWM}$ = 0.04, CrI = [0.00, 0.09],). Separate post-hoc models for low and high WM groups indicated that, whereas the drug slowed movement speed in the low WM group (Model 7.2: ($E\mu_{PLAvsHAL}$ = -0.08, CrI = [-0.16, -0.01], $P(E\mu_{PLAvsHAL} < 1)$ = 0.99), there were no drug effects on movement in the high WM group (Model 7.3: ($E\mu_{PLAvsHAL}$ = 0.00, CrI = [-0.04, 0.06]).

**Supplementary Table 7.1**. *Model parameters for model 7.1*

| Population-level effects | Estimate | Error | 95% CrI (lower) | 95% CrI (upper) |
|---|---|---|---|---|
| *Intercept* | 1.09 | 0.03 | 1.04 | 1.14 |
| *HAL vs PLA* | -0.04 | 0.02 | -0.08 | 0.01 |
| *Low WM* | -0.04 | 0.03 | -0.09 | 0.01 |
| *High WM* | 0.04 | 0.03 | -0.01 | 0.09 |
| *HAL vs PLA, low WM* | -0.04 | 0.02 | -0.09 | 0.00 |
| *HAL vs PLA, high WM* | 0.04 | 0.02 | 0.00 | 0.09 |
| **Group-level effects** | **Estimate (SD)** | **Error** | **95% CrI (lower)** | **95% CrI (upper)** |
| *Subject ID (Intercept)* | 0.14 | 0.02 | 0.11 | 0.18 |

*Note.* Model formula: speed ~ drug * WM + (1 | subject ID).

**Supplementary Table 7.2**. *Model parameters for model 7.2 (post-hoc model - low WM)*

| Population-level effects | Estimate | Error | 95% CrI (lower) | 95% CrI (upper) |
|---|---|---|---|---|
| *Intercept* | 1.05 | 0.04 | 0.97 | 1.13 |
| *HAL vs PLA* | -0.08 | 0.04 | -0.16 | -0.01 |
| **Group-level effects** | **Estimate (SD)** | **Error** | **95% CrI (lower)** | **95% CrI (upper)** |
| *Subject ID (Intercept)* | 0.12 | 0.04 | 0.04 | 0.20 |

*Note.* Model formula: speed ~ drug * WM + (1 | subject ID).

**Supplementary Table 7.3**. *Model parameters for model 7.3 (post-hoc model - high WM)*

| **Population-level effects** | **Estimate** | **Error** | **95% CrI (lower)** | **95% CrI (upper)** |
|---|---|---|---|---|
| *Intercept* | 1.13 | 0.04 | 1.05 | 1.22 |
| *HAL vs PLA* | 0.00 | 0.02 | -0.04 | 0.06 |

| **Group-level effects** | **Estimate (SD)** | **Error** | **95% CrI (lower)** | **95% CrI (upper)** |
|---|---|---|---|---|
| *Subject ID (Intercept)* | 0.15 | 0.03 | 0.10 | 0.23 |

*Note.* Model formula: speed ~ drug * WM + (1 | subject ID).

1 Vehtari, A., Gelman, A. & Gabry, J. Practical Bayesian model evaluation using leave-one-out cross-validation and WAIC. *Statistics and Computing* **27**, 1413-1432, doi:10.1007/s11222-016-9696-4 (2017).
2 Bürkner, P.-C. brms: An R Package for Bayesian Multilevel Models Using Stan. *2017* **80**, 28, doi:10.18637/jss.v080.i01 (2017).
3 rootSolve: Nonlinear root finding, equilibrium and steady-state analysis of ordinary differential equations (2009).
4 Cools, R., Gibbs, S. E., Miyakawa, A., Jagust, W. & D'Esposito, M. Working memory capacity predicts dopamine synthesis capacity in the human striatum. *J Neurosci* **28**, 1208-1212, doi:10.1523/jneurosci.4475-07.2008 (2008).
5 Landau, S. M., Lal, R., O'Neil, J. P., Baker, S. & Jagust, W. J. Striatal Dopamine and Working Memory. *Cerebral Cortex* **19**, 445-454, doi:10.1093/cercor/bhn095 (2009).
6 Schuster, B. A. *et al.* Dopaminergic Modulation of Dynamic Emotion Perception. *The Journal of Neuroscience* **42**, 4394, doi:10.1523/JNEUROSCI.2364-21.2022 (2022).
7 Kimberg, D. Y., D'Esposito, M. & Farah, M. J. Effects of bromocriptine on human subjects depend on working memory capacity. *NeuroReport* **8** (1997).
8 Mattay, V. S. *et al.* Effects of Dextroamphetamine on Cognitive Performance and Cortical Activation. *NeuroImage* **12**, 268-275, doi:10.1006/nimg.2000.0610 (2000).
9 Gibbs, S. E. & D'Esposito, M. Individual capacity differences predict working memory performance and prefrontal activity following dopamine receptor stimulation. *Cogn Affect Behav Neurosci* **5**, 212-221, doi:10.3758/cabn.5.2.212 (2005).
10 Frank, M. J. & O'Reilly, R. C. A mechanistic account of striatal dopamine function in human cognition: psychopharmacological studies with cabergoline and haloperidol. *Behav Neurosci* **120**, 497-517, doi:10.1037/0735-7044.120.3.497 (2006).
11 Rostami Kandroodi, M. *et al.* Effects of methylphenidate on reinforcement learning depend on working memory capacity. *Psychopharmacology* **238**, 3569-3584, doi:10.1007/s00213-021-05974-w (2021).
